# Efferent signaling along nociceptive peripheral terminals *in vivo* is enhanced during inflammation

**DOI:** 10.1101/2025.02.01.636018

**Authors:** Devora Gershon, Omer Barkai, Nurit Engelmayer, Ben Katz, Shaya Lev, Alexander M Binshtok

## Abstract

Primary nociceptors are essentially characterized as afferent neurons carrying noxious sensory information from the periphery to the CNS. However, the information flow on primary nociceptors is bidirectional. Nociceptor peripheral terminals release a variety of mediators to the target organ in the vicinity of the injured area. These mediators promote sensitization of adjacent sensory neurons, vasodilation, and edema and affect innate and adaptive immunity, leading to hyperalgesia and inflammation that often expands beyond the injured areas. Many theories associate these phenomena with the antidromic action potential propagation along nociceptor terminals; however, the antidromic efferent signaling at the single nociceptor terminals has never been demonstrated. Here, using *in vivo* calcium imaging from the individual nociceptive terminals innervating the mouse cornea together with a computational approach, we demonstrated that short-lasting activation of a single terminal *in vivo* was sufficient to activate the remote, non-activated terminal, which branches from the same nociceptor fiber. This increase was dependent on the activation of voltage-gated sodium and calcium channels. Moreover, we showed that the efferent signaling along nociceptive terminals increases under inflammatory conditions, culminating in enhanced calcium signaling in the remote non-activated terminals. This inflammation-induced increase in intra-terminal calcium could trigger the enhanced release of inflammatory mediators, spilling over wider areas and affecting terminals from adjacent unstimulated receptive fields, leading to the expansion of hyperalgesia and inflammation.

## Introduction

Primary nociceptor neurons encode the modality, intensity, and location of noxious stimuli and transmit this information to the CNS. Although devoid of synaptic connections, primary nociceptive afferents are not simple signal conductors but complex molecular structures with intricate architecture that can alter the sensory information flowing to the CNS^1–4^. These alterations could result from the modulation of the excitable properties of primary nociceptors by various mediators released from the injured tissue^5^, glial cells^6^, or sensory neurons themselves^1,7^. The latter, inter-nociceptive paracrine effects could be induced in the dorsal root ganglion (DRG)^7^ and at the target tissue via the release of various neuropeptides from the nociceptor peripheral terminals^8^. Glutamate, substance P (SP), calcitonin gene-related peptide (CGRP), and other neuromodulators that are released from the peripheral nociceptive terminal endings activate and sensitize sensory terminals^9–13^, often even in adjacent uninjured tissues, thus increasing their responsiveness to the applied stimuli, thereby expanding the hyperalgesic area^14^. In addition, the mediators released from nociceptive terminals can trigger neurogenic inflammation, yet again expanding the affected areas^8,15–21^. The vasoactive effects of these mediators (specifically, CGRP) are considered the triggers in initiating migraine attacks^22^. Notably, the release of neuropeptides from peripheral nociceptive terminals is critical for mediating local inflammation, apoptosis of dermal macrophages activating hair growth, and, importantly, for triggering the activation of local immune responses^15,20,23–27^. Therefore, understanding the mechanism of the peripheral release of neuropeptides by nociceptor terminals and its modulation in pathological conditions is of utmost importance.

At the effector organs, nociceptive terminals compose convoluted terminal trees by converging with other terminal branches from the same fiber^2,28^ (see also **Figure 1A, B**). When tissue is exposed to noxious stimuli, activation of the nociceptor terminal leads to action potential (APs) generation at the terminal^29^ that propagates orthodromically toward the CNS. Theoretically, when APs reach a convergence point with non-activated terminals of the same terminal tree, they could invade these terminals and propagate antidromically to their terminal tip - a phenomenon known as axon reflex^8,20^. A resulting depolarization at the terminal tip could be sufficient to activate voltage-gated calcium channels (VGCC) and trigger a rapid and local release of neural mediators from the non-activated terminals^30–32^, which in turn may induce vasodilation, plasma extravasation and edema, also affecting nociceptive excitability and triggering the activation of immune responses^8,20^. This local ortho-to-antidromic signaling is biophysically sound and implied in the literature as a possible mechanism for the release of the mediators from primary nociceptors^15^; however, the propagation of signals from a single activated terminal to the remote non-activated terminal was never demonstrated at the level of a terminal tree of a single nociceptor fiber.

**Figure 1.**
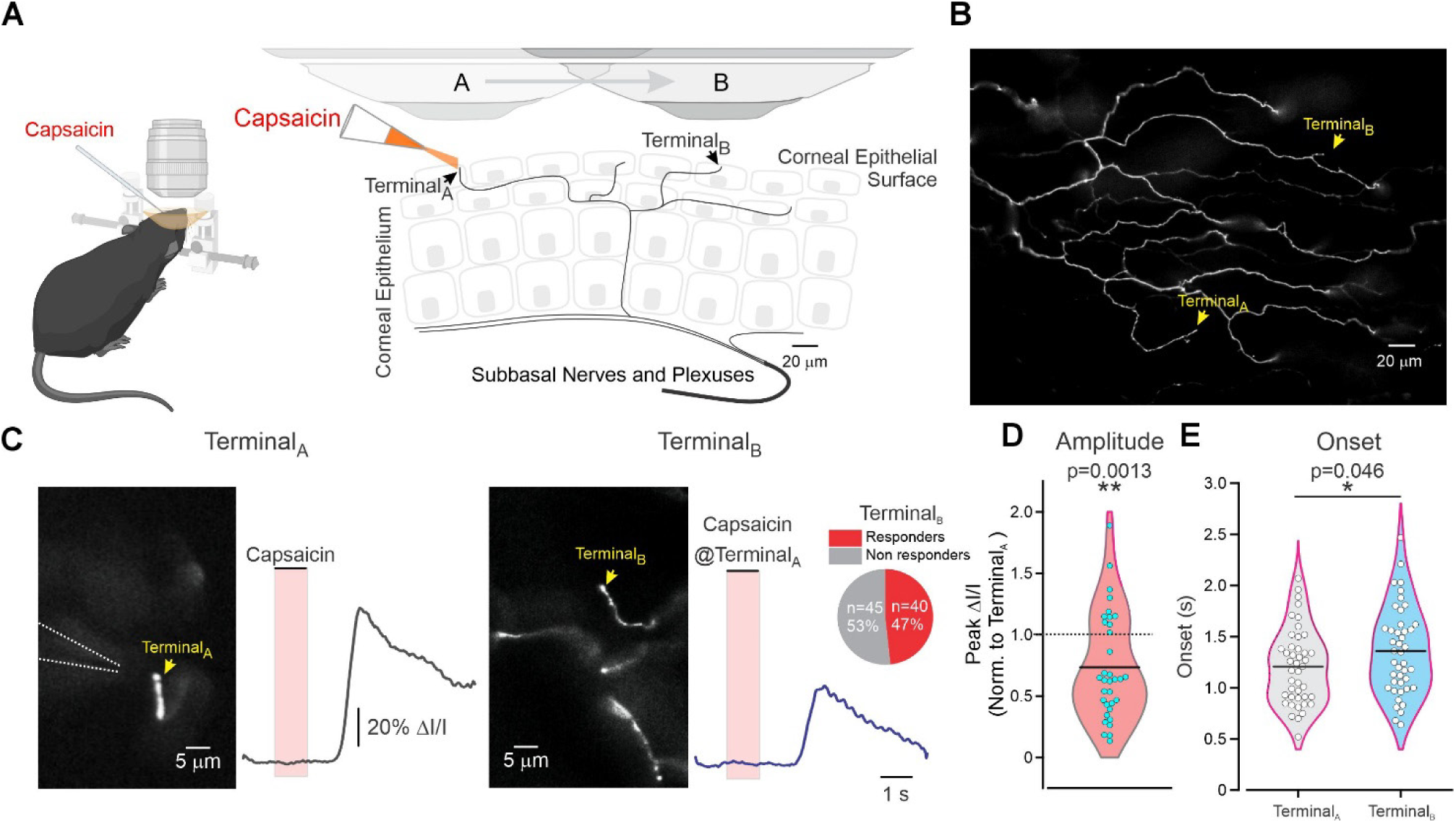
Activation of the nociceptor terminal triggers calcium elevation in the remote non-activated terminal of the same terminal tree. **A.** Scheme depicting the experimental procedure. First, changes in the intra-terminal calcium following focal calibrated puff application of capsaicin were recorded from the nociceptive terminal at the corneal superficial epithelium surface (Terminal_A_) in anesthetized, head-stabilized mice. Then, the lens was moved and refocused on a remote (at least 200 μm) terminal, which is a part of the same terminal tree, i.e., belongs to the same nociceptor neuron (Terminal_B_), but the puff pipette was kept at the same position adjacent to Terminal_A_. Five minutes after the recording from Terminal_A_, a second puff application to Terminal_A_ with the same parameters was applied, but the calcium responses were recorded from Terminal_B_. **B.** Representative image of a terminal tree with exemplary Terminal_A_ and Terminal_B_ (*arrows*). **C.** Representative epifluorescent images and traces of optical recording of the changes in fluorescent intensity from Terminal_A_ (*left*) and remote Terminal_B_ (*right*) following the application of 0.5 μM capsaicin on Terminal_A_. Representative of 40 out of 85 experiments. The yellow arrows indicate the region of interest (ROI) from which the recordings were performed. Dashed lines outline the location of the puff pipette for the activation of Terminal_A_. *Inset:* Pie chart of the fractions of Terminals_B_ that showed an increase in intra-terminal calcium (*responders, red*) or did not respond (*non-responders, gray*) to the activation of Terminal_A_ by capsaicin. n = 85 experiments from 85 eyes, N=85 mice. **D.** Violin plot showing the distribution of the ratios of peak fluorescence intensity measured in Terminal_B_ following activation of Terminal_A_, relative to the peak fluorescence intensity of Terminal_A_. The solid line represents the Median; the dotted line marks the ratio of “1”. Notably, most values fall below the ratio of “1”. Only terminal sets (Terminal_A_ and Terminal_B_) with successful Terminal_B_ responses were compared; one sample Wilcoxon signed rank test, n=40 terminals, N=40 mice. **E.** Violin plots comparing the Means (solid line) and the distribution of response onsets in Terminal_A_ and Terminal_B_ following the application of capsaicin onto Terminal_A_. Only terminal sets (Terminal_A_ and Terminal_B_) with successful Terminal_B_ responses were compared; paired *t*-test, n=40 terminals, N=40 mice.

Here, using *in vivo* calcium imaging of nociceptor terminals branching from a terminal tree of individual nociceptor neurons, we demonstrated that activation of a single terminal with noxious stimuli triggers an antidromic propagation along a single terminal tree, leading to an increase in calcium in non-activated terminals. This increase in calcium may underlie the release of mediators from nociceptor neurons and trigger the expanded hyperalgesia, neurogenic inflammation, and modulation of the immune responses. Moreover, we showed that efferent signaling along the terminals of a single terminal tree, and the calcium increase in the remote terminals, is enhanced under inflammatory conditions, potentially leading to elevated release of various mediators and altering the functionality of target organs.

## Results

### Axon reflex in nociceptor terminals: capsaicin-evoked activation of a specific terminal triggers activation of remote terminals of the same terminal tree

To demonstrate the existence of efferent signaling at the terminal tree of a single nociceptor fiber *in vivo*, we assessed the activity of unactivated individual nociceptor terminals following the activation of a single terminal of the same terminal tree. Accordingly, a combination of two Adeno-Associated Viruses (AAV) expressing RFP and GCaMPs was injected into the trigeminal nucleus (TG). Two weeks later, the expression of RFP and GCaMPs was observed *in vivo* in superficial ramified nociceptive terminals^33^ innervating the corneal epithelial surface (**Figure 1A, B**). First, we identified nociceptor terminal trees composed of multiple intraepithelial individual terminal branches and terminating in the superficial epithelium with a terminal tip (**Figure 1A, B**). We focally applied capsaicin (*see Methods*) on a specific terminal tip (activated terminal, Terminal_A_) and monitored the resulting changes in intra-terminal calcium (*see Methods*, **Figure 1A-C**). Next, we traced the axon of Terminal_A_ and its bifurcations along the terminal tree to a distant superficial terminal belonging to the same fiber (Terminal_B_, **Figure 1A, B**). To avoid the possibility of Terminal_B_ being activated due to capsaicin diffusion during Terminal_A_ activation, a distance of at least 200 μm between Terminal_A_ and Terminal_B_ was kept. This distance exceeds a previously determined calculated distance for capsaicin spillover^34^. While the lens was repositioned and refocused on Terminal_B_, the puff pipette was kept at the original position above Terminal_A_. After allowing full recovery of Terminal_A,_ which takes about 5 min^29^, a second puff application of capsaicin was given, and the calcium responses were recorded from Terminal_B_ (*see Methods*, **Figure 1A-C**).

We assumed that if efferent propagation exists, we should be able to detect an increase in calcium in Terminal_B_ following the activation of Terminal_A_. Indeed, we found that the activation of Terminal_A_ triggers calcium elevation in Terminal_B_ in ∼ 50% of cases (40 out of 85 terminals, 47%, **Figure 1C**, *inset*). The responses in Terminal_B_ activated following the activation of Terminal_A_ were significantly smaller than the responses in Terminal_A_ (Amplitude_TerminalB_/Amplitude_TerminalA_ = 0.74 ± 0.1, **Figure 1D**). As expected, the responses in Terminal_B_ following the activation of Terminal_A_ appeared with later onset (∼ 150 ms) than those in Terminal_A_ (**Figure 1E**).

Our previous computational data suggest that capsaicin-induced terminal depolarization in nociceptor-like terminals decays rapidly with distance in the absence of voltage-gated sodium channels (Na_V_) and the generation of action potentials (AP)^2^. Therefore, we hypothesized that the response in Terminal_B_ after the activation of Terminal_A_ is triggered by capsaicin-induced AP generation along Terminal_A_ that then propagates antidromically towards Terminal_B_. To examine this, we recorded the Terminal_A_-induced response of Terminal_B_ in the presence of the Na_V_ blocker oxybuprocaine^35^. Applying oxybuprocaine to mice corneas (*see Methods*) did not affect the capsaicin-induced activation of Terminal_A_ but completely and reversibly prevented calcium elevation in Terminal_B_ (**Figure 2**).

**Figure 2.**
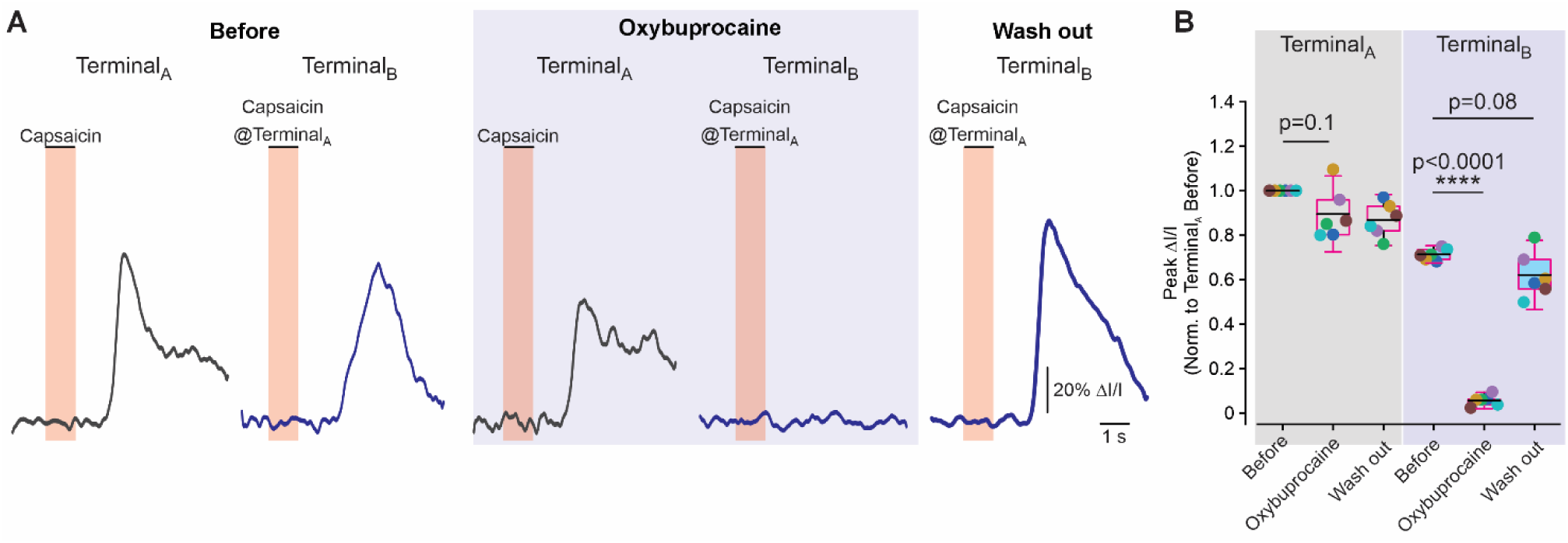
Blockade of voltage-gated sodium channels prevents activation of a remote terminal. **A.** Representative traces of optical recordings from Terminal_A_ (*left*) and Terminal_B_ (*right*) following activation of Terminal_A_ with 0.5 μM capsaicin before and after treatment with 0.4% oxybuprocaine and 1h after oxybuprocaine washout. Note that the response in Terminal_B_ was annulled after treatment with oxybuprocaine and restored following washout. Representative of 6 experiments. **B.** Box plots and individual paired values (color coded) of the fluorescence intensities of Terminal_A_ (*left*) and Terminal_B_ (*right*) following the application of capsaicin on Terminal_A_ before and after application of oxybuprocaine and after 1h of oxybuprocaine washout; RM one-way ANOVA with posthoc Bonferroni; n = 6 terminals from 6 different eyes from N=6 mice.

Altogether, these results demonstrate that activating a single nociceptor terminal triggers the activation of remote terminals, which requires antidromic AP propagation.

### Stimulation of the nociceptive terminal triggers activation of voltage-gated calcium channels at the remote unstimulated terminal

What could be the source of calcium increase in the remote Terminal_B_? The distance between Terminal_A_ and Terminal_B,_ which exceeds the calculated distance of capsaicin spillover and the dependence of calcium signals in Terminal_B_ on Na_V_-induced AP propagation, suggests that calcium elevation in Terminal_B_ is most likely independent of TRPV1 channel activation. Therefore, we hypothesized that calcium entry in remote terminals is mediated by voltage-gated calcium channels (VGCC) activated by depolarization from antidromically propagating APs. If the latter is the case, blockade of VGCC will affect Terminal_A_-induced calcium elevation in Terminal_B_. We and others demonstrated that nociceptor terminals express functional VGCC^29,36,37^ and that the inhibition of L-, N-, and T-type VGCC by benidipine^38^ abolishes electrically evoked calcium signals at the terminals^29^. These results suggest that L-, N-, and T-type VGCC constitutes the majority of VGCC at the terminals. Therefore, we examined the effect of blocking L-, N-, and T-type VGCC by benidipine on the responsiveness of Terminal_B_ following activation of Terminal_A_. Similar to our previous results^29^, the treatment with benidipine decreased the calcium response of Terminal_A_ to capsaicin (**Figure 3A**, *upper traces***, B**). Importantly, benidipine abolished the calcium signals in Terminal_B_ following Terminal_A_ activation (**Figure 3A**, *lower traces***, B**) in all terminals, including those where the capsaicin-induced signal was not, or only slightly, affected by benedictine (**Figure 3B**). These results suggest calcium entry at the remote non-activated terminal requires activation of VGCC.

**Figure 3.**
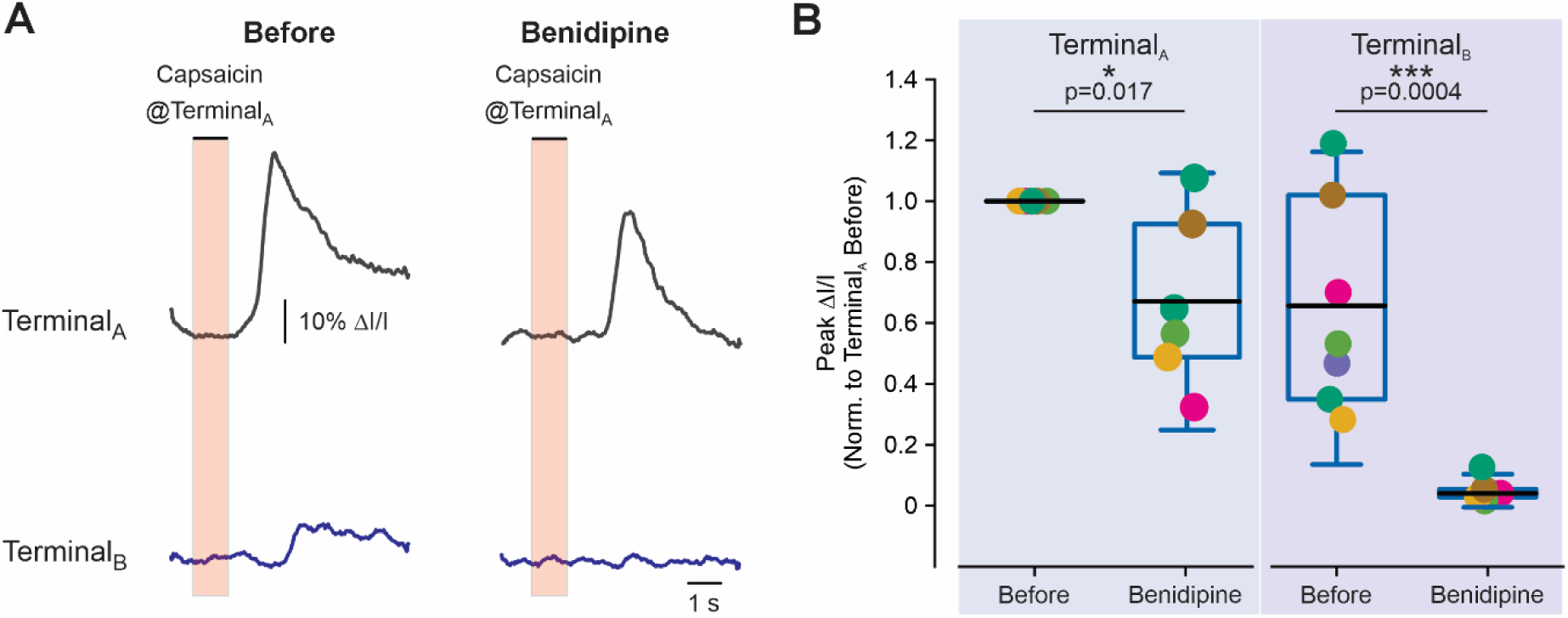
Blockade of voltage-gated calcium channels prevents calcium increase in a remote terminal. **A.** Representative traces of optical recordings from Terminal_A_ (*top*) and Terminal_B_ (*bottom*) following activation of Terminal_A_ with 0.5 μM capsaicin before and after 60 min incubation with benidipine (50 μM). **B.** Box plots and individual paired values (color coded) of the terminal fluorescence intensities of Terminal_A_ (*left*) and Terminal_B_ (*right*) following the application of capsaicin on Terminal_A_ before and after incubation with benidipine. Note that the application of benidipine prevents the activation of the remote terminals. One-sample t-test; n = 7 terminals from 7 different eyes from N=7 mice

### In simulated inflammation-like conditions, activation of the nociceptor terminal leads to increased antidromic propagation

Inflammation causes nociceptor hyperexcitability^5^, which could lead to an increase in both afferent and, consequently, efferent signaling along the terminal tree, thus amplifying the activation of remote terminals. We used a numerical model of nociceptor neurons with the terminal tree^2,29^ to predict how the efferent signaling changes in inflammatory conditions. To reflect our previous experimental findings, showing no functional availability of voltage-gated sodium channels (Na_V_) at the distal terminal part^29^, we annulled Na_V_ conductance along the first 25 μm of the terminal fiber such that the capsaicin-induced depolarization propagates passively over the first 25 μm of the terminal (Na_V_-less compartment). After 25 μm, a “propagation” part begins with a spike initiation zone (SIZ) in which normal sodium channel conductance is present^29^ (*see Methods*, **Figure 4A**). Under these control conditions, the activation of Terminal_A_ with simulated capsaicin-like current resulted in three depolarizing voltage deflections of ∼60 mV at the tip of Terminal_B_ (**Figure 4B**, *blue electrode and traces*). Interestingly, at the “propagation” zone of Terminal_B_, where voltage-gated sodium channels were present, the resulting depolarizatory deflections were ∼20 mV higher than those measured at the Terminal_B_ tip (ΔVm_Tip_ = 61.4±0.4 mV vs. ΔVm_SIZ_ = 83.7±2.8 mV, **Figure 4B**, *green electrode and traces*), suggesting a substantial voltage decay along with the “passive” Na_V_-less distal part of Terminal_B_.

**Figure 4.**
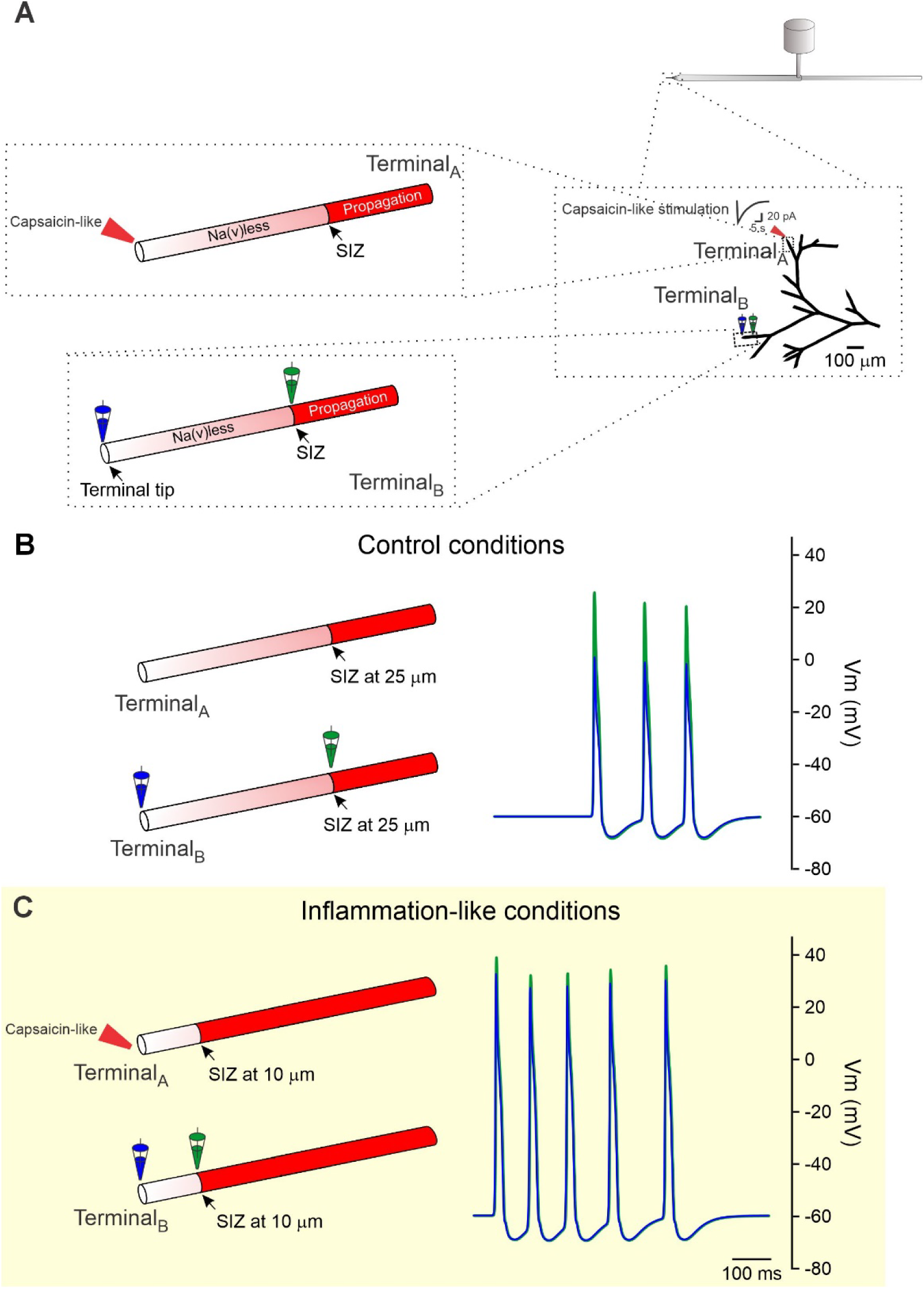
In simulated inflammatory-like conditions, the efferent signaling along the nociceptor terminal tree increases. **A.** A scheme depicting a modeled structure of nociceptive terminals with a terminal tree. Each terminal is composed of a distal compartment that does not express voltage-gated sodium channels (Na_V_-less, *grey*), which is limited by a spike initiation zone (SIZ) from which a compartment that contains Na_V_ conductance begins (“propagation” compartment, *red*). Terminal_A_ (indicated by the red pipette) was stimulated by a simulated capsaicin-like current (depicted near Terminal_A_), and the response was recorded from the tip (Na_V_-less, indicated by a blue electrode) and the “propagation” zone (green electrode) of Terminal_B_. **B.** *Left*, A scheme of a simulated control condition, in which an action potential is initiated in the SIZ of Terminal_A_, located 25 μm from the terminal tip. *Right*, overlapping traces recorded from the Terminal_B_’s tip (*blue*) and the “propagation” zone (*green*) following the activation of Terminal_A_ by a capsaicin-like current. Note a substantial decay of the depolarizing deflections from the “propagation” zone to the terminal tip. **C.** Same as *B,* but in simulated inflammatory-like conditions. In these conditions, the SIZ of Terminal_A_ and Terminal_B_ were shifted towards the terminal tip and located 10 μm from the tip. Note that in these conditions, similar activation of Terminal_A_ led to a more substantial depolarization at the “propagation” zone of Termnimal_B_ that only slightly decayed when it reached the tip of Terminal_B_.

Next, we simulated the inflammatory conditions by increasing the availability of Na_V_ at the terminal end, as we previously demonstrated^29^ (*see Methods*). We shifted the location of the SIZ towards the terminal tip so that the Na_V_-less zone ends 10 μm from the tip, and then the “propagation” zone begins. In these simulated inflammation-like conditions, applying the same capsaicin-like stimulation to Terminal_A_ generated five larger (∼90 mV) depolarizing voltage deflections in Terminal_B_ (**Figure 4C**, *blue electrode and traces*). The deflections detected at the Terminal_B_ tip were ∼5 mV lower than the responses recorded from the “propagation” zone (ΔVm_Tip_ = 90.2±2 mV vs. ΔVm_SIZ_ = 95.5±2.9 mV, **Figure 4C**, *green electrode and traces*).

These results predict that activation of the nociceptor terminal induces antidromic propagation. In inflammatory-like conditions, the antidromic propagation increases, leading to increased depolarization at the terminal tip partly due to a decreased decay in the depolarization that reaches the terminal end of the remote terminal.

### In inflammatory conditions, stimulation of the nociceptor terminal leads to increased activation of the remote terminals

Our computational results predicted that antidromic propagation towards unstimulated terminals increases under inflammatory conditions. Next, we examined whether, *in vivo,* under inflammatory conditions, the activation of a single terminal triggers a stronger activation of a remote terminal. Because of the high variability of the capsaicin-induced calcium responses due to the different depths of each terminal ending from the epithelial surface, a comparison can only be performed on the same terminal before and during inflammation (*see Methods*). Consequently, conventional inflammatory models that require comparisons of the terminal responses between different groups of animals were not applicable. Therefore, to examine the changes in efferent signaling in inflammatory conditions, we treated mice corneas with a combination of proinflammatory cytokines IL-1β and TNFα. We previously demonstrated that the combination of both cytokines leads to an increased response of the activated terminal to capsaicin^29^. Before examining the effect of the treatment on efferent signaling, we examined whether the application of IL-1β and TNFα is sufficient to trigger inflammatory hyperalgesia in mice. Indeed, applying capsaicin to the cornea after treatment with IL-1β and TNFα for 30 min significantly increased nocifensive behavior (**Figure 5A**). We therefore considered treatment with IL-1β and TNFα as an inflammatory model condition. Under these conditions, the responsiveness of Terminal_A_ to capsaicin increased, and activation of Terminal_A_ led to a significantly stronger response at Terminal_B_ (**Figure 5B, C**), which appeared with a shorter onset than in the control conditions (**Figure 5D**).

**Figure 5.**
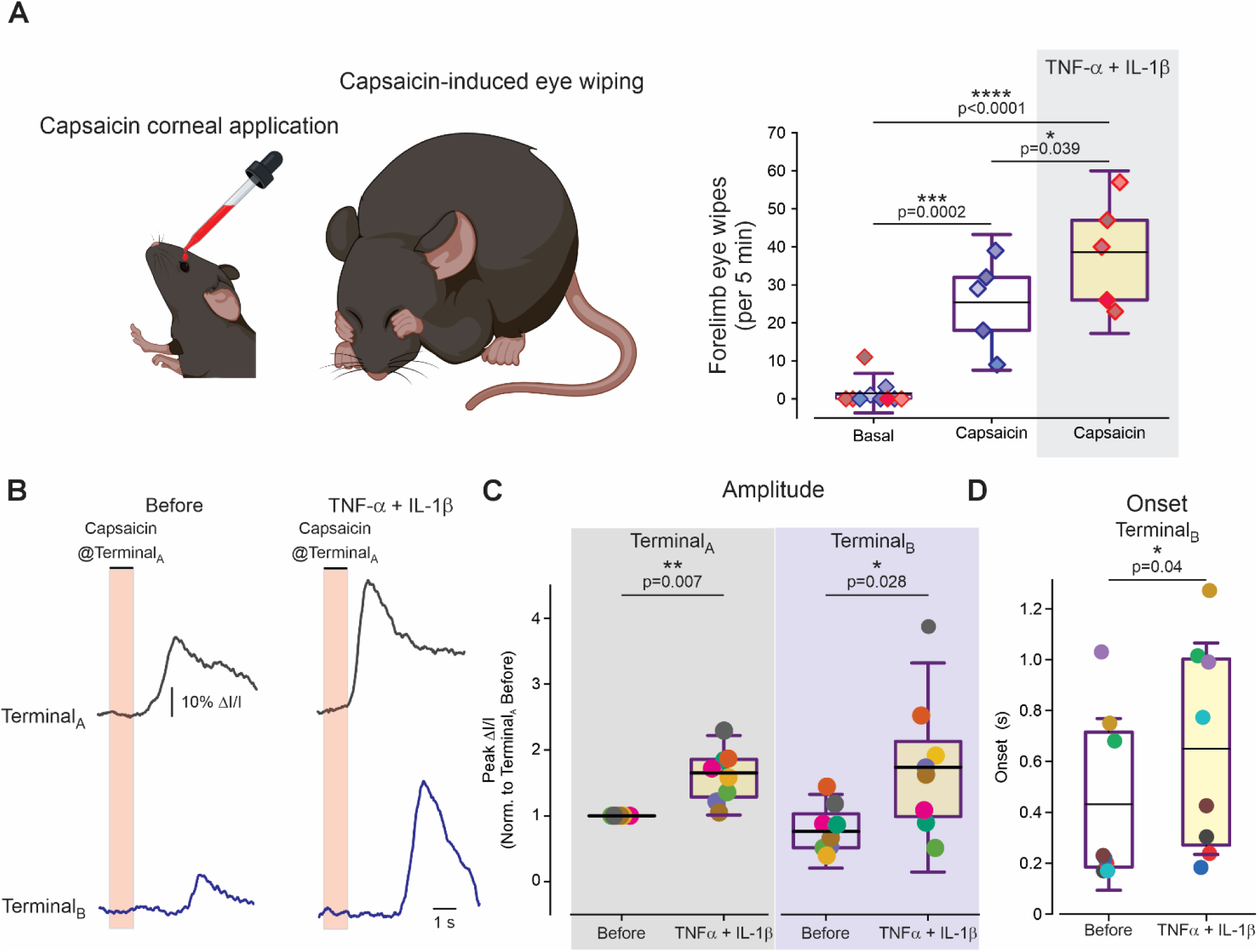
In the inflammatory conditions, activation of the nociceptor terminal triggers enhanced activation of a remote terminal. **A.** *Left*, scheme depicting a behavioral experiment. *Right*, Box plot and individual paired values (color coded) of the number of forelimb eye wipes after corneal application of saline (Basal), compared to 0.5 μM of capsaicin applied to mice pretreated with Vehicle or a combination of 100 ng/mL TNFα and 100 pg/mL IL-1β for 30 min (*see Methods*). Because no statistical differences were detected between mice in the Control group (p=0.55, unpaired *t*-test, N=10 mice), their data were pulled into one group (Basal), and ordinary one-way ANOVA with posthoc Bonferroni was performed to assess the statistical significance (N=5 mice in each group). One-way ANOVA performed when the Basal groups were separated according to the treatment shows similar statistical significance (p=0.04, comparing the effect of capsaicin applied to the vehicle and TNFα and IL-1β treated group, N=5 mice in each group). **B.** Representative traces of optical recordings from Terminal_A_ (*top*) and Terminal_B_ (*bottom*) following activation of Terminal_A_ with 0.5 μM capsaicin before and after 30 min co-incubation with 100 ng/mL TNFα and 100 pg/mL IL-1β. **C.** Box plots and individual paired values (color coded) of the terminal fluorescence intensities of Terminal_A_ (*left*) and Terminal_B_ (*right*), following the application of capsaicin on Terminal_A_ before and after incubation with TNFα and IL-1β. One-sample t-test; n = 8 terminals from 8 different eyes from N=8 mice. **D.** Box plots and individual paired values (color coded) of the normalized response onsets in Terminal_B_ following the activatiuoTerminal_A_ (before and after incubation with TNFα and IL-1β. The response onsets in Terminal_B_ were normalized to (subtracted from) the corresponding response onsets in Terminal_A_. Paired *t*-test; n = 8 terminals from 8 different eyes from N=8 mice.

## Discussion

The expansion of hyperalgesia and inflammation beyond the immediate injured area is well-described^39^. The axon reflex - an antidromic propagation of the APs along non-activated branches of the nociceptor that triggers the release of active substances from nociceptor neurons affecting sensory and immune systems - is thought to be responsible for this expansion^15,20^. However, although the axon reflex was predicted about 120 years ago^21^, antidromic propagation along a single nociceptive tree has not been demonstrated. Several alternative and additional mechanisms to explain the expansion of hyperalgesia and inflammation were suggested, including the activation of other nociceptor neurons via ephaptic connections between activated and non-activated axons and communication at the level of the sensory ganglion or dorsal horn via dorsal root reflexes^1,8^. Here, we utilized *in vivo* imaging from individual capsaicin-sensitive nociceptor terminals branching from the same terminal fiber and showed that a single activation of one nociceptor terminal with a short-lasting (1 s) application of noxious stimulus is sufficient to trigger an increase in calcium in a remote non-activated terminal of the same nociceptor neuron **(Figure 6A**, *left*).

**Figure 6.**
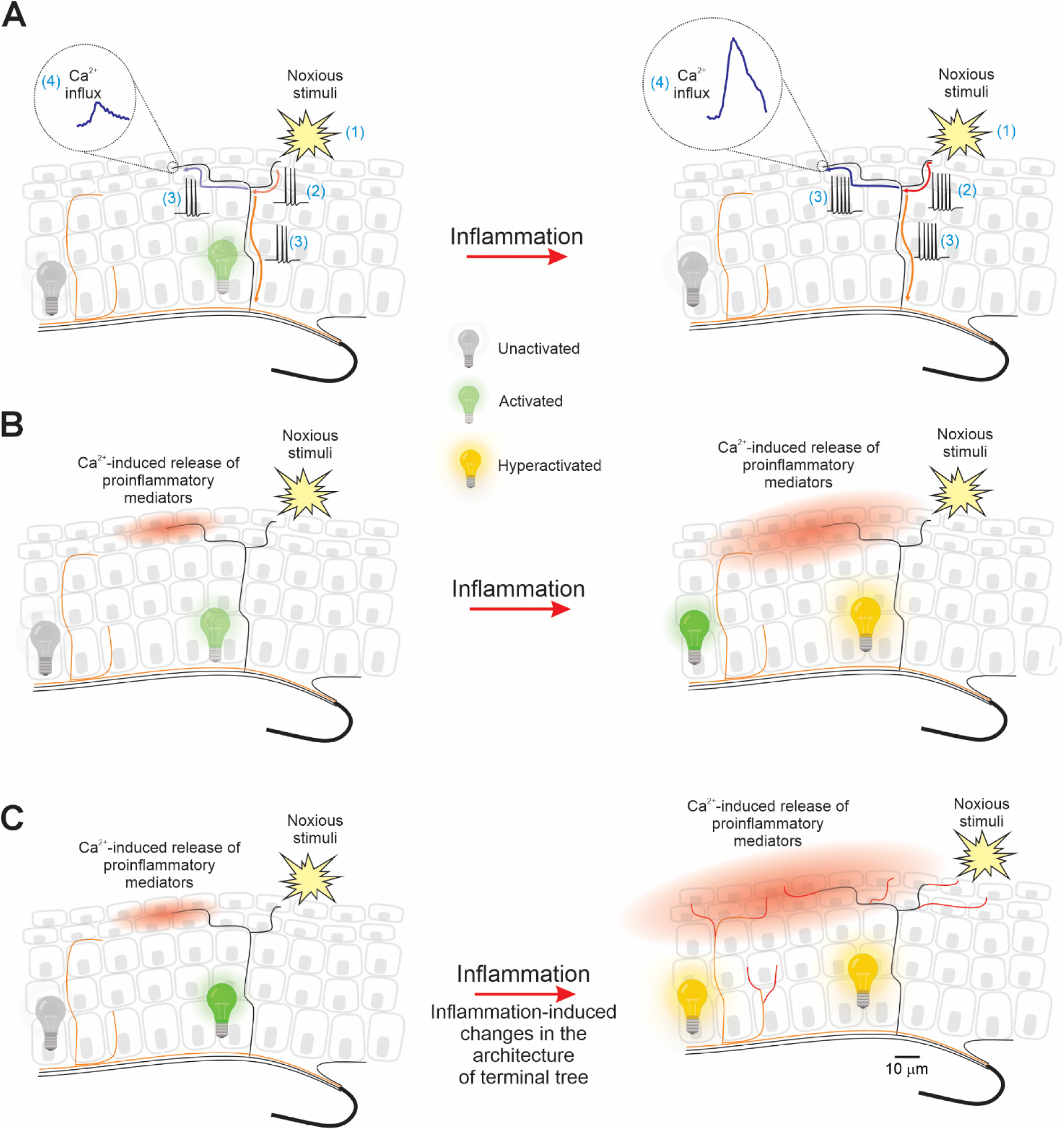
Scheme summarizing nociceptor axon reflex in normal and pathological conditions (A) and its potential implications on the expansion of hyperalgesia and neurogenic inflammation (B-C). **A.** *Left,* Activation of a specific nociceptor terminal (1) leads to the generation of APs (2), which propagates (3) both orthodromically (*orange arrow*) towards the CNS (“activated” *green lightbulb)* and antidromically (*blue arrow*) towards remote non-activated terminals of the same terminal tree. There, APs lead to an increase in intra-terminal calcium (4) via the activation of VGCC. *Right*, the antidromic signaling increases in inflammatory conditions, leading to an increased calcium signal at the remote terminal. The terminals of the nociceptors that innervate different receptive fields, which were not exposed to noxious stimuli, remained unactivated *(grey lightbulb)*. **B**, *Left*, In normal conditions, an axon reflex-induced increase in the intra-terminal calcium in remote terminals of peptidergic nociceptors triggers the release of various proinflammatory mediators (*red-shaded cloud*) that disperse around the remote terminal, resulting in neurogenic inflammation. *Right*, the inflammation leads to hyperexcitability of the activated terminal (“hyperactivated,” *yellow lightbulb*), also resulting in the inflammation-induced increase in intra-terminal calcium, which could result in the enhanced release of proinflammatory mediators, possibly reaching and affecting unactivated terminals of the nociceptors, innervating different receptive fields (*green lightbulb*), thus leading to the extension of hyperalgesia. **C.** Inflammation triggers structural changes in the nociceptor terminals^4^, characterized by the elongation of the individual terminal branches, thus increasing the possibility of activating the adjacent nociceptor fibers.

Our data suggest that this “remotely activated” calcium increase is mediated by the activation of VGCC following Na_V_-induced AP firing that propagates from an activated terminal. It has been demonstrated that the release of mediators from nociceptor terminals depends on calcium entry via VGCC ^31^, which initiates the SNARE-dependent release of neuropeptides^30,32^. Given that our study focused exclusively on capsaicin-sensitive (TRPV1-expressing) neurons, which are primarily peptidergic ^40^, it is reasonable to suggest that VGCC-dependent calcium increase in the remote terminal we described here may trigger the release of nociceptive mediators^41^ (**Figure 6B**, *left*). Importantly, we demonstrated that under inflammatory conditions sufficient to produce hyperalgesia, activating the nociceptor terminal leads to an enhanced calcium increase in the remote terminal **(Figure 6A**, *right*). In theory, this increased activation could, in turn, result in an increased release of mediators from the remote terminal (**Figure 6B**, *right*), leading to their extensive spillover and further increasing neurogenic inflammation. Additionally, the released mediators can reach terminals of nociceptors and other sensory fibers innervating adjacent areas (**Figure 6B**, *right*), increasing their excitability^9,12,13^, thus leading to the expansion of hyperalgesia beyond the injured area. Recent reports of inflammation-induced increases in nociceptor terminal branching and the length of each terminal branch^4^, together with the enhanced efferent release of nociceptive mediators from the terminal branches, may further contribute to the expansion of their effects on more expansive areas (**Figure 6C**).

Our previous results demonstrated that the nociceptive terminal is composed of two compartments: the “Na_V_-less” compartment, devoid of functional Na_V_s in which the signals propagate passively, and the axonal or “propagation” compartment, where Na_V_s are functionally available and signals propagate in the form of AP^29,34^. At the border between these two compartments, about 25 μm from the terminal tip, the spike initiation zone (SIZ, the area where AP is generated) is located^29^. Thus, APs invading the remote terminals would not regenerate after reaching the SIZ (because there are no functionally available Na_V_s from the SIZ until the tip), and the depolarization would decay as it passively propagates further towards the terminal tip (**Figure 4B**). This decay may result from the small diameter of the terminal branch and intra-terminal mitochondria^42–44^, which increases terminal axial resistance and decreases the length constant^2^. This voltage decay could affect the activation of VGCC and, consequently, the release of the nociceptive mediators. In inflammatory conditions and, in particular, following the treatment with TNFα and IL-1β, the increase in the responsiveness of the nociceptive terminals was attributed to the increase in the availability of the Na_V_s at the terminals, leading to a shift of the SIZ towards the terminal tip^29^. Assuming that during inflammation, the shift in SIZ and the increase of the available Na_V_s along the terminal tip occurs in all terminal branches, AP that propagates autonomically could invade the remote terminal closer to the terminal tip, leading to a smaller decay of the depolarization reaching the tip, as our computational model predicts (**Figure 4C**). This could be one of the reasons for the inflammation-induced increase in calcium signaling in remote terminals. Our data, which demonstrate a decrease in the onset time of the response of the remote terminal following the application of capsaicin on the activated terminal in the inflammatory conditions, also support this hypothesis.

We demonstrated that the axon reflex appears only in 50% of the remote terminals. This is puzzling, considering our data showing AP-mediated propagation between activated and remote terminals, which implies that the signal will not decay on its way to the remote terminal. However, as shown in Figure 4 and discussed above, the signal decays along with the passive Na_V_-less part of the terminal. The axial resistance of this part may vary among different terminals due to the differences in terminal diameter and number of intra-terminal mitochondria^42–44^. Thus, in the terminals with smaller diameters or larger amounts of intra-terminal mitochondria, the voltage decay between the “propagation” part and the passive terminal tip could be strong enough to prevent the activation of VGCC, thus precluding calcium increase in the remote terminals. Another possibility is that in some cases, capsaicin-mediated calcium influx induced by activation of TRPV1 and VGCC in Terminal_A_ fails to generate APs at the Terminal_A_ SIZs.

Our experiments demonstrating calcium increase in the remote terminals following the application of capsaicin on the activated terminal show the efferent signaling in nociceptor terminals; however, they do not directly prove that this signaling resulted from the antidromic AP propagation. Our computational data predicts the ortho-to-antidromic AP propagation at the level of the branches of the terminal tree (see also Barlai et al.^2^). We previously demonstrated that applying Na_V_ blocker oxybuprocaine prevents the increase in calcium in the terminal fibers proximally to the SIZ^29^, suggesting the propagation beyond this point is AP-dependent. Together with the data showing that a blockade of Na_V_ prevents efferent signaling, it suggests local antidromic AP transmission towards non-activated terminals. However, recent findings demonstrating that nociceptive neurons have an axon initial segment (AIS) near their somata^45^ may suggest that the efferent signaling we observe *in vivo* may also occur following AP propagating orthodromically and generating AP at the somatic AIS, which then propagates antidromically, activating the terminals of the same nociceptor neuron.

We show that applying noxious stimuli on a nociceptor terminal evokes calcium signals in the remote terminals that depend on the activation of N-, L-, and/or T-type VGCC. We previously demonstrated that functional L-, T-, or N-type VGCC are expressed at the terminal tip^29^. Blockade of N-and L-type VGCC has been shown to prevent the electrical stimulus-evoked increase in CGRP in the skin *in vitro*, suggesting the involvement of these channels in neuropeptide release^31^. T-type VGCC that is expressed by nociceptor peripheral terminal processes^37,46^ was also related to the evoked release of CGRP from nociceptor neurons^47^. Altogether, these results suggest that L-, N- and T-type-mediated calcium increase following efferent signaling could trigger the release of nociceptive mediators from the remote terminals.

It is widely accepted that inflammation increases nociceptor excitability^5,48–50^, leading to enhanced responses of the nociceptor terminals to the noxious stimuli^29^, plausibly increasing antidromic propagation and consequently efferent signaling. We modeled inflammatory conditions *in vivo* by treating cornea with a combination of proinflammatory cytokines TNFα and IL-1β, which, as we and others previously demonstrated, are released during inflammation^51–55^ and acutely increase AP firing in nociceptor neurons^56–58^. Moreover, we showed that the short-lasting application of TNFα and IL-1β on mouse cornea *in vivo* increases the capsaicin-induced calcium response in nociceptive terminals^29^. However, treating corneas with TNFα and IL-1β does not fully recapitulate the inflammatory state. We used this approach, despite its limitation, because it allowed us to directly compare the changes in the efferent signaling in the same terminal before and after the induction of the acute inflammation. Our data demonstrating that similar short-lasting TNFα and IL-1β treatment that led to an increase in efferent signaling was sufficient to induce corneal hyperalgesia partly overcame the limitation of the model. However, we cannot exclude that the alterations of the efferent signaling in the “natural” inflammatory state could differ from those following exposure to proinflammatory cytokines.

In summary, our results provide evidence for the existence of the axon reflex at the level of individual nociceptor terminal trees and its enhancement in inflammatory conditions. We show that acute activation of the nociceptor terminal triggers calcium elevation at the remote, non-activated terminals, which, if sufficient, could induce the release of nociceptor mediators. The calcium elevation and plausibly mediator release is increased in inflammatory conditions. Considering the role of the release of nociceptor mediators in migraine, inflammation, and coordinated integration with the immune cells, our results may advance our understanding of the mechanistic and temporal aspects of these interactions, providing a basis for better therapies for pain, inflammation, and immunopathologies.

## Resource Availability

All data needed to evaluate the conclusions are presented in the paper. All the data and the materials are fully available upon request from the lead contact. All model parameters and the complete code used for simulation are available on the ModelDB repository (Accession: 266850).

## Acknowledgments

This work was funded by: The Israel Science Foundation – Individual research grant 1202/23 (AB), Israel Cancer Research Fund (ICRF) – The Brause Family Initiative for Quality of Life 22-402-QOL (AB), The Israel Science Foundation (ISF) and the Azrieli Foundation - 2545/18 (AB), and Cecile and Seymour Alpert Chair in Pain Research (AB).

## Author Contributions

Conceptualization, A.B.; Investigation, D.G., O.B., N.E. and S.L.; Formal Analysis, D.G., O.B., N.E., B.K., S.L. and A.B.; Writing – Original Draft, A.B.; Writing – Review & Editing, B.K., and A.B.; Funding Acquisition, A.B.; Supervision, A.B.

**Declaration of interests**

The authors declare that they have no competing interests.

## Metherials and Methods

### Animals

Adult (4-6 weeks, 20-25 gr) male C57BL/C mice were used, and all procedures were approved by the Ethics Committee of the Hebrew University (Ethic protocol number MD-21-16546-4 and MD-18-15608) in accordance with the institutional and national guidelines and regulations.

Animals were housed in a specific pathogen-free (SPF) environment in standard husbandry and housing conditions, according to the regulations of the Hebrew University of Jerusalem Authority of Biological and Biomedical Models. Specifically, animals were housed in groups under a controlled temperature (23 ± 2°C) and humidity-controlled environment, with ad libitum access to food and water. Animals were kept in a 12-h light/dark cycle in standard cages with bedding and environment enrichment. Animals were randomly allocated and assigned to experimental groups.

### Viral vectors

Adeno-Associated Virus serotype 1 (AAV1) pENN.AAV.hSyn.TurboRFP.WPRE.RBG deposited by James M. Wilson (Addgene plasmid # 105552; http://n2t.net/addgene:105552; RRID: Addgene_105552) expressing a TurboRFP gene, and Adeno-Associated Virus serotype 1 (AAV1) pAAV.Syn.GCaMP6s.WPRE.SV40 deposited by Douglas Kim & GENIE Project (Addgene plasmid # 100843; http://n2t.net/addgene:100843; RRID: Addgene_100843) expressing a calcium sensor GCaMP6, were ordered from Addgene, delivered on dry ice, and stored at –80°C.

A mixture of the two viruses at a ratio of 1:3 of RFP to GCaMP6s (both with a titer of ≥ 1×10¹³ vg/mL) was made and aliquoted for a working solution of 9 µl.

### Viral injection

The expression of the calcium and RFP indicators at the corneal terminals was achieved by viral infection of the trigeminal ganglion (TG) cell bodies via stereotaxic injection to the V1 area as we previously detailed^29,59^. In short, an adult mouse was anesthetized by intraperitoneal (IP) injection of 100 µl of prepared anesthesia mixture containing ketamine (1 gr/10 ml) and medetomidine (1 mg/ml) diluted in saline (NaCl 0.9%) up to 10 ml, to final concentrations of 10 mg/ml ketamine and 80 µg/ml medetomidine. For pain relief, meloxicam (5 mg/ml) was injected subcutaneously at a concentration of 1.32-1.65 mg/kg. The mouse’s scalp was shaved, and the mouse was placed in the stereotactic frame on a heating pad set at 37°C. The mouse was anesthetized using an isoflurane vaporizer (0.4%-0.6%) connected via a nose cone. 2% of isoflurane was set for anesthesia induction, and 0.4-0.6% was used to maintain the anesthesia during the procedure.

Animals were monitored for pain reflexes using the paw pinch method. When reflexes were abolished, the animal’s head was stabilized using non-rupture ear bars, and depilatory cream was applied to remove the remaining fur after shaving. The scalp was cleaned using saline and 70% ethanol, followed by applying 10% povidone-iodine. A midline incision of about 1 cm was made to expose the skull to visualize Bregma and Lambda focal points used for coordinates determination.

The coordinates for cranial drilling allowing the viral injection into the V1 area of the two trigeminal ganglions (TGs) were adopted from Whitehead et al.^60^: on the mediolateral axis, two holes (approximately 0.5 mm diameter) were drilled according to the following coordinates: +0.4 mm ± 0.02 mm to Bregma, on the anterior-posterior axis, and +/–1.35 ± 0.02 mm to Bregma on the mediolateral axis. After drilling a hole, the dental drill was replaced with a pipette holder for the viral injections. A calibrated 1-5 μl glass pipette loaded with the viral mixture of RFP and GCaMP6 was mounted into the holder. 1-2 μl of the viral mixture was injected per ganglion at a rate of about 1 µl/10 s. After injection of the virus, the incision was closed using a suture thread, and the animal was injected IP with atipamezole (5 mg/ml) diluted in saline to a final concentration of 1-1.25 mg/kg for anesthesia reversal. The calcium imaging experiments from the terminals of corneal nociceptive nerve endings were performed 10 - 14 days after the injections.

### *In vivo* calcium imaging of corneal nociceptive terminals

#### Experimental design

For a step-by-step explanation of the *in vivo* calcium imaging from corneal nociceptive terminals, please refer to Gershon et al.^59^

The signal propagation between the activated terminal (Terminal_A_) and the remote, non-activated terminal (Terminal_B_) of the same corneal nociceptive terminal tree was assessed using the following approach (**Figure 1A**): mice infected with calcium (GCaMP6s) and fluorescent (RFP) indicators (see above *Virus injection* section) were anesthetized with a combination of ketamine and medetomidine by IP injection and placed on a plate with a heating pad set to 37°C. We used ketamine (1 gr/10 ml) and medetomidine (1 mg/ml) diluted in saline (NaCl 0.9%) up to 10 ml, to final concentrations of 10 mg/ml ketamine and 80 µg/ml medetomidine. When the paw pinch reflex was abolished, the head was fixed with a three-point head stabilizer (SGM-4, Narishige, Japan). A custom-made eye stabilizer and bath was used to stabilize the eyeball and filled with standard extracellular solution (SES) composed of (in mM): 145 mM NaCl, 5 mM KCl, 2 mM CaCl_2_, 1 mM MgCl_2_, 10 mM glucose, and 10 mM HEPES. The eye bath was held by a hemostat fixed to a Noga arm. The plate was placed on the microscope stage. Using RFP fluorescence (see below *Optics and fluorescence* section for the parameters), a superficial (less than 5 μm from the epithelial surface) ramified nociceptive terminal was selected (Terminal_A_), and its coordinates were stored. Then, by shifting the microscope lens, its bifurcations were followed until a distant superficial terminal belonging to the same tree was located (Terminal_B_). A minimal distance of 200 μm between Terminal_A_ and Terminal_B_ were kept to avoid the effect of capsaicin spillover. The experiment began by activating Terminal_A_ with a calibrated capsaicin puff application (see below, *Terminal activation* section). Wide-field epifluorescence and calcium imaging recordings were performed to record the induced calcium signals at the Terminal_A_ 2-10 μm proximal to the terminal tip ^59^. Then, the lens was moved and refocused on Terminal_B_, but the puff pipette was kept at the same position adjacent to Terminal_A_. Five minutes after the recording from Terminal_A_, a second puff application to Terminal_A_ with the same parameters was given, but the calcium responses were recorded from Terminal_B_ (**Figure 1A-C**). We previously demonstrated that the 5-minute interval between the stimulations allows a full recovery of the terminal from desensitization^29^.

Only the experiments showing a successful Terminal_B_ activation were taken for the analysis.

#### Optics and fluorescence parameters

Wide-field epifluorescence and calcium dynamics imaging were performed using an Olympus BX51WI microscope with an x40 LUMPlanFL objective and an NA of 0.8. An Exi Aqua monochromatic camera (QImaging) was used for wide-field epifluorescence image acquisition controlled by NIS Elements AR software Version 4.20.02, Nikon software. The RFP fluorescence imaging was performed using an exposure time of 300-500 ms, binning 2X2, hardware gain of 10-20, and a maximal power light source with 3.1 mW/cm^2^ flux at the focal plane of the objective. A back-illuminated 80 X 80 pixel cooled CCD camera (NeuroCCD-SMQ, RedShirt Imaging) was used for fast optical recording of changes in GCaMP6s fluorescent intensity, set at a 40-125 Hz acquisition rate and medium gain. Fluorescent excitation was performed with a CoolLed fluorescence excitation system. For RFP, a 565 nm excitation LED and RFP filter set (Ex 560, Em 630, dichroic Lp 585; Chroma) were used. For GCaMP6s, a 490 nm excitation LED (maximal power light source with 5.4 mW/cm^2^ flux at the focal plane of the objective) and GFP filter set (Ex 480, Em 535, dichroic Lp 510; Chroma) were used.

The ROIs were visualized using a NeuroCCD-SMQ camera and Turbo-SM recording software. Turbo-SM via the RedShirt camera’s analog-to-digital (A/D) converter, triggering a Digidata 1440 A/D interface (Molecular devices), which in turn triggers the picospritzer (Pneumatic PicoPump) was used to initiate the puff and start the Redshirt camera recordings. The experimental parameters were set using the pClamp software (Molecular Devices). These parameters have been devised after calibrating the dispersion profile of the puffed solution (Goldstein et al., 2017), preventing an effect on neighboring terminals. The experimental protocol was as follows: 1000 ms recording before puff application, 1000 ms puff duration (2 pounds per square inch (psi)), and a 5500 ms recording following the puff.

Image data collected from the RedShirt camera was further processed and analyzed from selected ROIs (defined with a kernel size of two (2X2 pixels) using NeuroPlex 10.2.0, MATLAB software (Mathworks, version R2016a), and OriginPro 2020 (OriginLab Corporation). Using the NeuroPlex program provided by RedShirt, the raw files of the ROI fluorescent intensities recorded were assessed. The data was further analyzed by MATLAB, the fluorescent traces were normalized to baseline fluorescent intensities (I_0_), and the changes in the peak fluorescence intensity (I/I_0_) and signal time onset (in ms) were calculated^29,59^.

#### Terminal activation

The recordings from Terminal_A_ and Terminal_B_ were performed following a focal activation of Terminal_A_ by capsaicin puff. To that end, a pulled glass pipette (see ^59^ for specifications) was filled with a solution of SES, capsaicin (500 nM) and sulfa-rhodamine 101 (SR101, 9 μM) (4-6 MΩ resistance) and was placed in the Picospritzer holder adapter on the micromanipulator (SM7) (UNIT Junior RE, Luigs & Neumann). The loaded pipette tip was positioned adjacent to the terminal of interest, 2-5 μm above the corneal surface (without disturbing the cell layer), and approximately 10 μm in the x-y plane from the terminal end, and 1 s puff was applied^29,59^.

To examine the effect of voltage-gated sodium channels (Na_V_) blockade on efferent signaling, a 10μl drop containing 0.4% oxybuprocaine in standard external solution (SES) was applied to the eye bath for 2 min, and the responses to capsaicin were recorded. Afterward, the solution was washed out and replaced with SES. 60 min later, the effects of washing out oxybuprocaine were assessed. Only terminals showing successful Terminal_B_ activation before the application of oxybuprocaine were taken for the analysis.

To examine the effect of voltage-gated calcium channels (VGCC) blockade on efferent signaling, the efferent signaling was measured before and after 60 min pretreatment with 50 μM of the L-N- and T-type VGCC blocker benidipine (Alomone, Israel). A 10 mM stock solution was made up of ethanol and diluted to 50 μM in SES for working bath application. Only terminals showing successful Terminal_B_ activation before the application of benidipine were taken for the analysis.

#### Induction of corneal inflammation in vivo

Corneal inflammation was induced using a 10μl drop containing 100ng/ml of TNFα and 100pg/ml of IL-1β in SES applied to the eye every 5 min for a total duration of 30 min. Subsequently, the solution was washed out and replaced with SES for imaging. Calcium imaging of the same terminal was performed before and after the inflammation.

### Behavioral experiments

To ensure that the application of 100ng/ml of TNFα and 100pg/ml of IL-1β produces hyperalgesia, capsaicin-induced nocifensive behavior was analyzed using the forelimb eye wiping test as described previously^4,59,61^. Briefly, 6 male C57BL/6 mice (6-8 weeks old) were used in each experimental group. Before all experiments, mice were handled and habituated to the experimental procedures and testing environment for 4 days. The behavioral assays were performed in custom Perspex glass cages. The mice were habituated to the setup for 10 minutes before each assay, and then baseline eye-wiping behavior was recorded for 5 minutes. Following this, the animals were divided into experimental groups. The mice in the “Inflammation” group were anesthetized using isoflurane, and then a 10μl drop containing 100ng/ml of TNFα and 100pg/ml of IL-1β in standard external solution (SES) was applied to the right eye every 5 min for a total duration of 30 min. After 30 min, the animals were removed from anesthesia, and the behavioral essay began. The animals in the “Vehicle” group underwent the same procedure, but only SES was applied. The SES consisted (in mM): 145 mM NaCl, 5 mM KCl, 2 mM CaCl_2_, 1 mM MgCl_2_, 10 mM glucose, and 10 mM HEPES.

Each assay involved applying a 10μl drop containing 1μM capsaicin on the right eye of the mouse. Five minutes after application, the eye-wiping behavior was recorded and then analyzed post-hoc.

### Computational model

The model was built based on a published, open-source biophysical model of unmyelinated axons in DRG neurons ^2,4,29^. The model files are accessible in the ModelDB repository (Accession: 266850). Simulations were conducted using the NEURON simulation environment. The nociceptor morphology included a soma-like compartment with a 25 μm diameter connected to a stem axon that branched into central and peripheral axons meeting at a T-junction. Branch diameters were 0.25 μm.

### Passive Membrane Properties

Intrinsic membrane properties were adapted from Barkai et al.^2^. A passive membrane resistance of 10,000 Ω·cm² was applied to all compartments except the terminal branch. Axial resistance was set to 150 Ω·cm for all compartments except the terminal branch, which exhibited a 4-fold somatic membrane resistance. The membrane capacitance for all compartments was 1 μF·cm², and the passive reversal potential (EPas) was set to −60 mV.

### Active Conductances

The model incorporated several active conductances, including TTX-sensitive sodium currents (INaTTXS), TTX-sensitive persistent sodium currents (INaP), Nav1.9 TTX-resistant sodium channels (INav1.9), and Nav1.8 TTX-resistant sodium channels (INav1.8). Parameters for these sodium channels were adapted from Herzog et al. and Baker et al.^62,63^. Three potassium channel types were included: (i) the delayed rectifier potassium channel (IKDR), adapted from Herzog et al.^62^; (ii) an A-type potassium channel (IKA) based on Miyasho et al.^64^ with activation and inactivation gates shifted by 20 mV in the hyperpolarized direction to match DRG neuron kinetics^65^; and (iii) Kv7/M channels adapted from Shah et al. ^66^ with activation curve parameters tuned as in Barkai et al. ^67^. The h-current (Ih) was included based on Shah et al. (Shah et al., 2002), with the slope factor adjusted according to Komagiri and Kitamura^68^. T-type (ICaT) and L-type (ICaL) calcium channels represented low voltage-activated (LVA) and high voltage-activated (HVA) currents, respectively. Unless otherwise noted, specific channel conductances (g) were as published previously^2–4:^ *g_Nav1.8_ = 0.02 S/cm*^2^*, g_Nav1.9_ = 0.00064 S/cm*^2^*, g_NaTTXS_ = 0.0017 S/cm*^2^*, g_NaP_ = 0.00005 S/cm*^2^*, g_KDR_ = 0.00083 S/cm*^2^*, g_KA_ = 0.0015 S/cm*^2^*, g_Kv7/M_ = 0.00034 S/cm*^2^*, g_H_ = 0.00033 S/cm*^2^*, g_CaL_ = 0.003 S/cm*^2^*, g_CaT_ = 0.001 S/cm*^2^.

### Reversal Potentials

The reversal potentials for sodium (ENa), potassium (EK), and the h (EH) currents were set as follows: ENa = 60 mV, EK = −85 mV, and EH = −20 mV. Sodium conductances were unevenly distributed, localized primarily in conductive compartments, while other conductances were uniformly distributed across all compartments.

Based on our previous findings, the terminal branches were divided into two sections separated by the spike initiation zone (SIZ)^29^. The SIZ consisted of a 25 μm-long compartment completely devoid of sodium conductances (Nav-less compartment). This Nav-less compartment was connected to the rest of the terminal branch via the "propagation" compartment, which bridged it to the junction^2,29^.

### Nerve-Ending Stimulation

Capsaicin-like stimulation was modeled and applied as in previous studies ^2–4,29,67^. A capsaicin-like current was introduced through a simplified voltage-clamp point process with fast exponential activation and slow exponential inactivation. The modeled capsaicin-like current mimicked the experimental kinetics of puff-applied 1 μM capsaicin-induced currents, sufficient to induce action potential (AP) firing in acutely dissociated DRG neurons^69^.

The axial resistance (Ra) of the terminal was increased by x15 relative to the distal axon Ra to simulate the localization of intracellular organelles in the free-ending terminal branches^2,44,70^.

Transducer channel conductance was introduced into the stimulated nerve endings to simulate the activation of transducer channels during capsaicin stimulation^2^. The conductance followed an exponential distribution, with the decay constant (γ) calculated to reflect capsaicin diffusion and concentration changes as a function of distance from the pipette tip, as determined in earlier work ^29^. Recordings were conducted by positioning a NEURON "point-process" electrode at the terminal end of the central axon ^2,67^.

To simulate the inflammatory conditions, the location of SIZ was shifted towards the terminal tip by shortening the Nav-less compartment by 15 μm, such that the SIZ was located 10 μm from the terminal tip ^29^.

### Statistical analysis

Data from the individual terminals is presented for each data set. We used one terminal set (Terminal_A_ and Terminal_B_) per mouse so that each data point represented individual mice. Due to possible variance in fluorescent intensities at different focal planes, only the values acquired at the same focal plane at the same ROI were compared. Fluorescent intensities were normalized to the first puff activation, and only paired statistical analysis was used to compare the fluorescent intensities at the same ROI before and after treatment. Because all the values were normalized to the values before treatment, a one-sample t-test or Wilcoxon signed rank test (when the values were not normally distributed) was used. To analyze the changes in response onsets between Terminal_A_ and Terminal_B,_ the unnormalized values were compared using paired *t*-test. The normality was assessed using the Shapiro-Wilk test. For the behavioral experiments, ordinary one-way ANOVA with posthoc Bonferroni was used. Actual p values are presented for each data set. The criterion for statistical significance was p < 0.05. Boxplots presented in the figures depict the Means or Medians (when the values were not normally distributed), 25th; 75th percentile, and 1.5 SD.

## Notes

### Competing Interest Statement

The authors have declared no competing interest.

